# *Kiss1* neurons in the medial amygdala of mice are sexually dimorphic and unique from hypothalamic anteroventral periventricular *Kiss1* neurons in their action potential firing properties

**DOI:** 10.64898/2026.09.19.752853

**Authors:** Leonie M. Pakulat, Yuanxin Zhang, Elisande Guillet, Madeline McGinnis, Maria Lesslie, William H. Colledge, Susan Jones

## Abstract

The importance of hypothalamic kisspeptin (*Kiss1*) neurons in controlling the reproductive axis is well established, while a population of extra-hypothalamic *Kiss1* neurons is found in the medial amygdala (MeA) of both males and females in rodents and is less well understood. The MeA is connected with the olfactory bulbs, the hypothalamus and cortical areas. MeA *Kiss1* neurons have been implicated in sexual behaviour and modulation of the reproductive axis; however, little is known about their action potential firing properties, which would influence kisspeptin release in target brain regions, nor whether their physiological properties are sex- or estrous cycle-dependent. In this study, whole-cell patch-clamp recordings were made from *Kiss1* neurons in brain slices containing the MeA of adult female (estrous and diestrous) and male mice. MeA *Kiss1* (*Kiss1^MeA^*) neurons were sexually dimorphic, with greater numbers in male compared to female mice and a slower change in excitability in estrous females compared with males. However, combining a range of electrophysiological parameters using principal component analysis did not reveal a distinct sex- or estrous cycle-linked phenotype, suggesting that the properties of *Kiss1^MeA^* neurons- and therefore kisspeptin release- are broadly similar. Interestingly, *Kiss1^MeA^* neurons had unique electrophysiological properties compared with anteroventral periventricular (AVPV) hypothalamic *Kiss1* (*Kiss1^AVPV^*) neurons in female mice; with their reduced excitability, achieving action potential firing frequencies greater than 10Hz required a larger current input, indicating that *Kiss1^MeA^* neurons require stronger synaptic input to achieve firing frequencies that drive kisspeptin release. These findings provide new information about the physiological properties of *Kiss1^MeA^*neurons, showing subtle sex differences and profound regional differences between the MeA and AVPV.

## Introduction

Kisspeptin is well established as the neuropeptide driving puberty and the reproductive axis by acting on gonadotropin-releasing hormone (*GnRH*) neurons in the preoptic area of the hypothalamus to stimulate luteinizing (LH) and follicle stimulating hormone (FSH) release from the pituitary^1–6^. This system is conserved between humans and rodents, with deletions in either the gene encoding kisspeptin (*Kiss1*) or its receptor (*Gpr54*/*Kiss1r*) causing hypogonadotropic hypogonadism across species^7,8^. The *Kiss1* and *Kiss1r* genes are expressed in brain regions beyond the reproductive hypothalamic areas in humans and rodents^9–15^. In humans, intravenous injection of kisspeptin enhances brain activity in the amygdala and other limbic brain regions in response to visual sexual and couple-bonding non-sexual stimuli, as well as a female olfactory stimulus^16,17^. Kisspeptin increases penile tumescence and self-reported arousal to sexual stimuli in men with hypoactive sexual desire disorder in a randomized clinical trial, suggesting a potential for therapeutic application^18^. In rodents, *Kiss1*^MeA^ neurons are implicated in modulating LH release in males and females, pubertal timing in females and partner preference in males^19–26^. The rodent MeA receives direct input from the accessory and main olfactory bulb and integrates important social olfactory information^27^. A functional connection between olfactory bulb *GnRH* neurons and *Kiss1*^MeA^ neurons, relevant for mounting during copulation, has been demonstrated in male mice^28^.

The physiological properties of *Kiss1*^MeA^ neurons are relatively unknown compared with hypothalamic *Kiss1*^AVPV^ neurons and *Kiss1* neurons in the arcuate nucleus (*Kiss1*^ARC^). *Kiss1*^AVPV^ neurons are sexually dimorphic in rodents, with greater numbers in females than males^12,13,29^, regulate the pre-ovulatory LH surge^30,31^, and are implicated in sexual behaviour (lordosis) in females^19,32–34^. *Kiss1*^ARC^ neurons are abundant in both sexes and regulate the pulsatile release of LH^35^. *Kiss1*^ARC^ and *Kiss1*^AVPV^ neuronal properties are well characterised and share fundamental similarities: they are relatively small with high input resistance and the ability to fire fast bursts of action potentials^36^. Neuronal firing in hypothalamic *Kiss1* neurons is modulated by sex steroids in intact and ovariectomized animals^36^. Like *Kiss1*^AVPV^ neurons, *Kiss1*^MeA^ neurons are sexually dimorphic, with greater numbers in male compared with female rodents^12,13,29^ and they are implicated in sexual behaviours. However, nothing is known about the electrophysiological properties of *Kiss1*^MeA^ neurons^36^. *Kiss1*^MeA^ neurons express genes encoding estrogen, androgen, and progesterone receptors and *Kiss1* transcription is upregulated by estrogen^37–40^, but it is not known how sex steroids modulate *Kiss1*^MeA^ neuronal properties. In this study, the action potential firing properties of *Kiss1*^MeA^ neurons was investigated using brain slices from adult male versus estrous and diestrous female mice, in order to determine sex- and estrous-cycle dependent effects on the activity of *Kiss1*^MeA^ neurons. *Kiss1*^MeA^ neuronal firing was also compared with *Kiss1*^AVPV^ neurons from female mice. *Kiss1*^MeA^ neurons show subtle sex differences in their firing properties and have a unique electrophysiological profile compared to *Kiss1*^AVPV^ neurons.

## Methods

### Animals

Mice were bred from heterozygous *Kiss1-Cre*^15^ and homozygous tdTomato mice (*Gt(ROSA)26Sor^tm9(CAG-tdTomato)Hze^*/J; stock #007909; The Jackson Laboratory) (Kiss1^tm2(Cre-GFP)Coll^-tdT), hereafter referred to as *Kiss1^Cre-tdT^*, to visualize *Kiss1* neurons. After the CRE-mediated recombination event tdT expression is independent of continued *Kiss1* promoter activity. Genotyping was performed as previously described^15^. For a comparison of tdTomato labelling in different *Kiss1*-Cre models, the same tdT reporter mice were crossed with Kiss1Cre(v2) mice, originally generated in the Palmiter Lab^41^ and kindly donated by Professor Allan Herbison; these are hereafter referred to as *Kiss1-Cre-v2-tdT*. Experimental animals were housed in open-top group cages with environmental enrichment on a reversed light-cycle with lights off at 7am and on at 7pm. Animals were sexually naïve unless otherwise specified. All experiments were conducted under the UK Home Office Animals (Scientific Procedures) Act 1986, following ethical review by the University of Cambridge.

### Quantification of tdTomato positive neurons

To quantify the number of tdTomato positive (tdT^+^) cells in the MeA, brain tissue was collected from adult mice of both sexes in both lines of mice at different ages throughout adulthood, as well as 19-20 week (wk) males with sexual experience (post-breeding), females at the time of the first vaginal plug (plugged) or after raising a minimum of one litter (post-breeding). Brains were postfixed in 4% paraformaldehyde/PBS overnight, washed in phosphate-buffered saline (PBS) and sectioned to 50 μm (coronal) using a Leica VT100S vibratome. All sections (anterior-posterior -0.8 to -2.8 coordinates) were imaged on an Evos FL fluorescent microscope with a 4x objective and all tdT+ positive cells lateral to the optic tract on each side were counted. Brains from one male and one female at 19 weeks old from the *Kiss1-Cre-v2-tdT* line were sectioned and imaged more extensively. Each coronal slice was visually assigned to coordinates in the Paxinos brain atlas^42^ by examining morphology.

### RNAscope in-situ hybridization

Male and female adult mice between 24 and 31 weeks were killed by cervical dislocation and brains quickly extracted, flash frozen in isopentane on dry ice and stored at -80 until sectioning. Sections (12μm) containing both the MeA and the ARC were cut on a cryostat. In situ hybridization for *Kiss1* (Catalog No. 500141, Advanced Cell Diagnostics (ACD), USA) and *tdTomato* (Catalog No. 317041-C2, ACD, USA) was performed using the RNAscope 2.5 HD Duplex Assay (Catalog No. 322430, ACD, USA), with pre-treatment conditions for fresh-frozen tissue as specified by the manufacturer except that sections were pre-treated with Protease IV for 15 minutes, AMP5 incubation time was reduced to 15 minutes and red dye incubation time was reduced to 8 minutes. Images of the MeA and ARC were taken as RGB images on an Evos FL fluorescent microscope with a 40x objective. For analysis, nuclei were segmented in CellPose^43^ and red and green signal manually annotated in Qupath^44^.

### Immunohistochemistry

Immunohistochemistry was used to confirm that the tdTomato neurons in the MeA are *Kiss1* neurons by visualization of the CRE protein expressed from the *Kiss1-Cre* allele. Brains from *Kiss1^Cre-tdT^*mice were perfusion fixed (4% paraformaldehyde/PBS) and post-fixed overnight at 4°C and transferred to 30% sucrose/PBS at 4°C until they sank. Brains were frozen in OCT and 20 µm coronal sections cut through the brain region containing the MeA. Sections were air dried for 2h at room temperature, rehydrated in TBS (20 mM Tris, 150 mM NaCl, pH7.5), subjected to antigen retrieval (1mg/ml trypsin in H_2_0, 10 min, room-temperature) and blocked with TBS + 0.3% Triton X-100 (TBST), 0.3M glycine, 5% BSA (2h, room temperature). The slides were incubated overnight at room temperature with the anti-CRE primary antibody (Rabbit monoclonal D7L7L, Cell Signalling Technology, 1:500 diluted in TBST + 0.5% goat serum), washed in TBST and then incubated with an Alexa488 labelled goat anti-rabbit secondary antibody (Invitrogen UK, Cat # A-11008, 1:500 diluted in TBST + 0.5% goat serum) for 1h at room temperature. After washing, sections were mounted in Fluoromount G + DAPI (Catalog No 00-4959-52, Thermo Fisher Scientific, UK) and visualized either on an EVOS M5000 microscope (Thermo Fisher Scientific, UK) or by confocal microscopy (Carl Zeiss, LSM 900).

### Brain slice preparation for electrophysiology

All electrophysiology experiments were carried out in *Kiss1^Cre-tdT^*mice. Estrous cycle regularity in females was determined by cytological examination^46^ of vaginal lavage for a minimum of 10 days prior to electrophysiological recordings. On experimental days, vaginal lavage was performed ∼1-2 hours before tissue collection. Only females in estrous or diestrous were used for electrophysiological recordings, with estrous defined as a sample dominant in cornified cells preceded by proestrous cytology the previous day. Diestrous was defined as being preceded by an estrous sample two days prior with a metestrous sample the day in between.

Adult mice aged 13 to 22 weeks were killed by cervical dislocation (in the morning of the experiment, 2-4 hours after lights off) and the brain was immediately extracted and submerged in ice-cold solution (in mM: 206 sucrose, 2.5 KCl, 1.25 NaH_2_PO_4_, 26 NaHCO_3_, 20 glucose, 5 MgCl_2_, 1 CaCl_2_; pH 7.4 with 95% O_2_/5% CO_2_). Coronal brain slices (190-220μm) containing the MeA were cut on a Vibrating Microtome (7000smz; Campden Instruments, UK) and left to recover for a minimum of 1 hour in an incubation chamber containing (in mM): 119 NaCl, 2.5 KCl, 1.25 NaH_2_PO_4_, 26 NaHCO_3_, 20 glucose, 5 MgCl_2_, 2 CaCl_2_; pH 7.4 with 95% O_2_/5% CO_2_ at 30°C.

### Whole-cell patch-clamp recordings

Brain slices were perfused with solution containing (in mM): 119 NaCl, 2.5 KCl, 1.25 NaH_2_PO_4_, 26 NaHCO_3_, 20 glucose, 1 MgCl_2_, 2 CaCl_2_, pH 7.4 with 95% O_2_/5% CO_2_ at 30±2 °C (Warner Instruments). *Kiss1*^MeA^ and *Kiss1*^AVPV^ neurons were identified as tdT+ using an LED at 554nm; non-*Kiss1*^MeA^ neurons were chosen based on absence of fluorescent signal. Whole-cell patch-clamp recordings were made as described previously^45^. Patch pipettes were 2-5MΩ when filled with solution (in mM: 120 K^+^-Gluconate, 5 KCl, 3 MgCl_2_, 2.8 NaCl, 20 HEPES (4-(2-hydroxyethyl)-1-piperazineethanesulfonic acid), 2 Na_2_ATP, 0.2 Na_3_GTP, 0.5 CaCl_2_, 5 EGTA (Ethylene glycol-bis(2-aminoethylether)-*N,N,N′,N′*-tetra-acetic acid), pH 7.2-7.3, 270-285 milliosmoles). Series resistance (Rs) and input resistance (Ri) were measured throughout and recordings were excluded if Rs exceeded 30 MΩ or either Rs or Ri changed by more than 25%. At the start of each recording, resting membrane potential (RMP) was measured (values have been corrected for a liquid junction potential of -18mV). To record action potential properties, membrane potential was held at -70mV in current-clamp and a depolarising current pulse was administered for either 10 or 500ms in 5pA increments. To measure the hyperpolarization-activated time-dependent voltage sag (Vsag), hyperpolarizing current was injected for 3s. All data were recorded using an Axopatch 200B amplifier (Molecular Devices, USA), low pass filtered (2kHz) and acquired to Spike 2 software (Version 10; Cambridge Electronic Design, Cambridge, UK) via a Micro1401 at a sampling frequency of 20kHz.

### Whole-cell patch-clamp data analysis

Spike trains were analysed using Spike 2. Maximum spike number is the maximum number of spikes (crossing 0mV) during a 500ms current pulse; dv/dt of Current*Spike Number is the dv/dt of a linear regression fitted to the 10% minimum to 90% maximum of the current input-spike output curve; rheobase is the lowest current pulse at which an action potential was initiated; frequency was measured at the current pulse giving the maximum number of spikes; mean coefficient of variation was measured across all current pulses; first AP delay was measured from current pulse onset to the peak of the first spike at rheobase; medium afterhyperpolarization (mAHP) was measured as the difference between the post-train trough and the starting voltage (within 300ms after the 500ms pulse) at rheobase; spike frequency adaptation (SFA) was calculated as the percentage of the last divided by the first inter-spike-interval at the current pulse evoking the maximum number of spikes (ISI last/first)%; depolarization block was measured as the decrease of equal/more than 20% from the maximum in the number of spikes at the highest (160pA) current pulse. Voltage sag was measured as the difference between the most hyperpolarized voltage and the steady state voltage in response to a 3s hyperpolarizing current pulse.

Single action potential shape parameters were analysed in DataView (version 11.21.4; Dr. W. J. Heitler, University of St Andrews). Action potential (AP) threshold level was determined by a change in inflection in a phase plot of the AP; AP amplitude is the difference between the threshold and AP peak; AP width was measured at the half height of the AP (WHH); the fast afterhyperpolarization (fAHP) was measured as the difference between the starting voltage and the hyperpolarization occurring within 3ms of the AP peak; the rise rate is the maximum dv/dt of the rising phase of the AP from threshold to peak and the fall rate is the maximum dv/dt of the falling phase of the AP from peak to trough.

### Statistical Analysis

All statistical analyses was performed using GraphPad Prism v10 and data represented as mean ± standard error (SEM). Normality was assessed with the Shapiro-Wilk normality test followed by a one-way ANOVA with Tukey’s post hoc test, a Brown-Forsythe and Welch ANOVA, or a Kruskal Wallis (with Dunn’s post hoc test) where appropriate. Data was standardized to a mean of 0 and standard deviation of 1 for principal component analysis (PCA), where principal components (PCs) are based on parallel analysis.

## Results

### Kiss1-tdTomato labelling in the MeA increases throughout adulthood in both sexes

To examine the number of *Kiss1* expressing neurons in the MeA of gonadally intact *Kiss1^Cre-tdT^* mice, tdTomato fluorescence was used since *Kiss1*-driven CRE recombination activates the tdTomato reporter gene. Representative sections through the MeA of male and female *Kiss1^Cre-tdT^* mice are shown in Figure 1A; the number of tdT^+^ cells in the MeA significantly increased throughout adulthood in both sexes. Male mice showed higher numbers of tdT^+^ cells than females at all equivalent ages (Figure 1B, C). The anterior-posterior range in which tdT^+^ cells were found was similar in both sexes (AP: -0.95 to -2.45) (Figure 1D). Male mice showed a peak number of tdT^+^ cells at posterior coordinates while in females the peak of tdT^+^ cells was more anterior. To investigate whether more cells in the MeA might be recruited to express *Kiss1* in response to reproductive events, the tdT^+^ cell count was examined in females and males (19-20 wks old) following mating. There was no effect of sexual experience on the number of tdT^+^ cells in males or females (Figure E, F).

**Figure 1.**
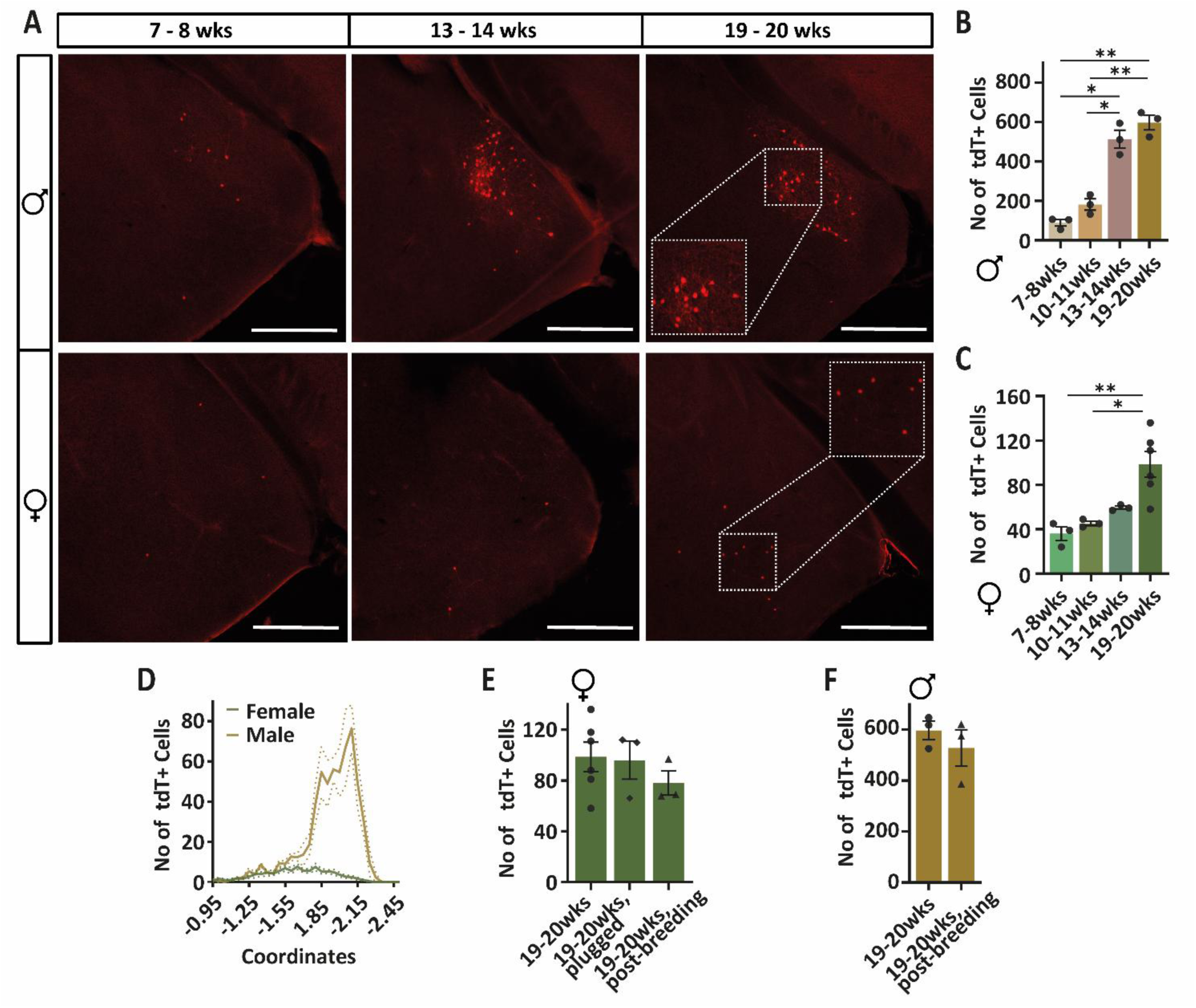
TdT labelling in the MeA is sexually dimorphic and increases with age. A. Representative images of tdT^+^ cell distribution in 50µm slices (Scale bar = 500µm). B, C. Number of tdT^+^ cells at 7-8, 10-11, 13 -14, and 19-20 wk old in male and female *Kiss1^Cre-tdT^* mice (F: Brown-Forsythe ANOVA F(3, 6.9) = 17.81, *p* = 0.0013; M: One-Way ANOVA F = 53.82, *p* < 0.0001). D. Mean anterior to posterior distribution of tdT^+^ cells in both sexes at 19-20 wks. E. Number of tdT^+^ cells of 19-20 wk old female mice culled with different sexual/breeding experience (virgin, day of the first vaginal plug, after one or multiple litters reared until weaning) (One-Way ANOVA F (2, 9) = 0.6829, *p* = 0.53). F. Number of tdT^+^ cells of 19-20 wk old virgin and sexually experienced male mice (Welch’s t-test t(2.99) = 0.86, *p* = 0.455). All post hoc P values: \**p* < 0.05; \*\**p* < 0.01. Data from 3 mice in all cases except 19-20wk females (6 mice).

As the number of tdT^+^ neurons in the *Kiss1^Cre-tdT^*females was relatively low, for comparison we examined the number of TdT^+^ neurons in the widely used *Kiss1^Cre-v2-tdT^* line (Figure S1). Both males and females of the *Kiss1^Cre-v2-tdT^* line had more tdT^+^ cells (Figure S1 A, B) but they showed a similar pattern of spatial distribution and temporal expression (Figure S1 C). We also observed tdT^+^ cells in brain areas not usually associated with *Kiss1* expression in the *Kiss1^Cre-v2-tdT^* line (Figure S2).

### MeA tdTomato positive neurons in Kiss1Cre-tdT mice express the Kiss1 gene and the CRE protein

To confirm that the tdT^+^ neurons in the *Kiss1^Cre-tdT^*mice still express the *Kiss1* gene, chromogenic dual in-situ hybridization (RNAscope) for *tdTomato* and *Kiss1* transcripts was performed. *Kiss1* expression in tdT^+^ cells in the ARC has been well documented in this transgenic mouse line; therefore, the ARC was used as a positive control in this study (Figure 2A). Very few *tdT*^+^ neurons could be visualized in the MeA of female mice due to sparse labelling in the thin sections and so only males were examined for MeA *Kiss1* expression. Around 50% of tdT^+^ cells contained more than 3 *Kiss1* puncta; this was not significantly different from the 60% co-expressing in the ARC (Figure 2B). In both regions, there were some neurons that expressed *Kiss1* but not *tdT*, this occurred more often in the MeA (Figure 2A,C). There were no differences in the amount of *Kiss1* puncta per neuron between *Kiss1*^MeA^ and *Kiss1*^ARC^. As an additional confirmation of whether tdT^+^ cells are actively transcribing *Kiss1*, immunohistochemistry for CRE was performed in male *Kiss1^Cre-tdT^*mice. In this mouse model CRE is only produced when the *Kiss1* promoter is active. In the MeA, 87 ± 2.6% (n=3 mice) of tdT^+^ cells were immunoreactive for CRE and 89 ± 3.9% (n=5 mice) in the ARC (Figure2D).

**Figure 2.**
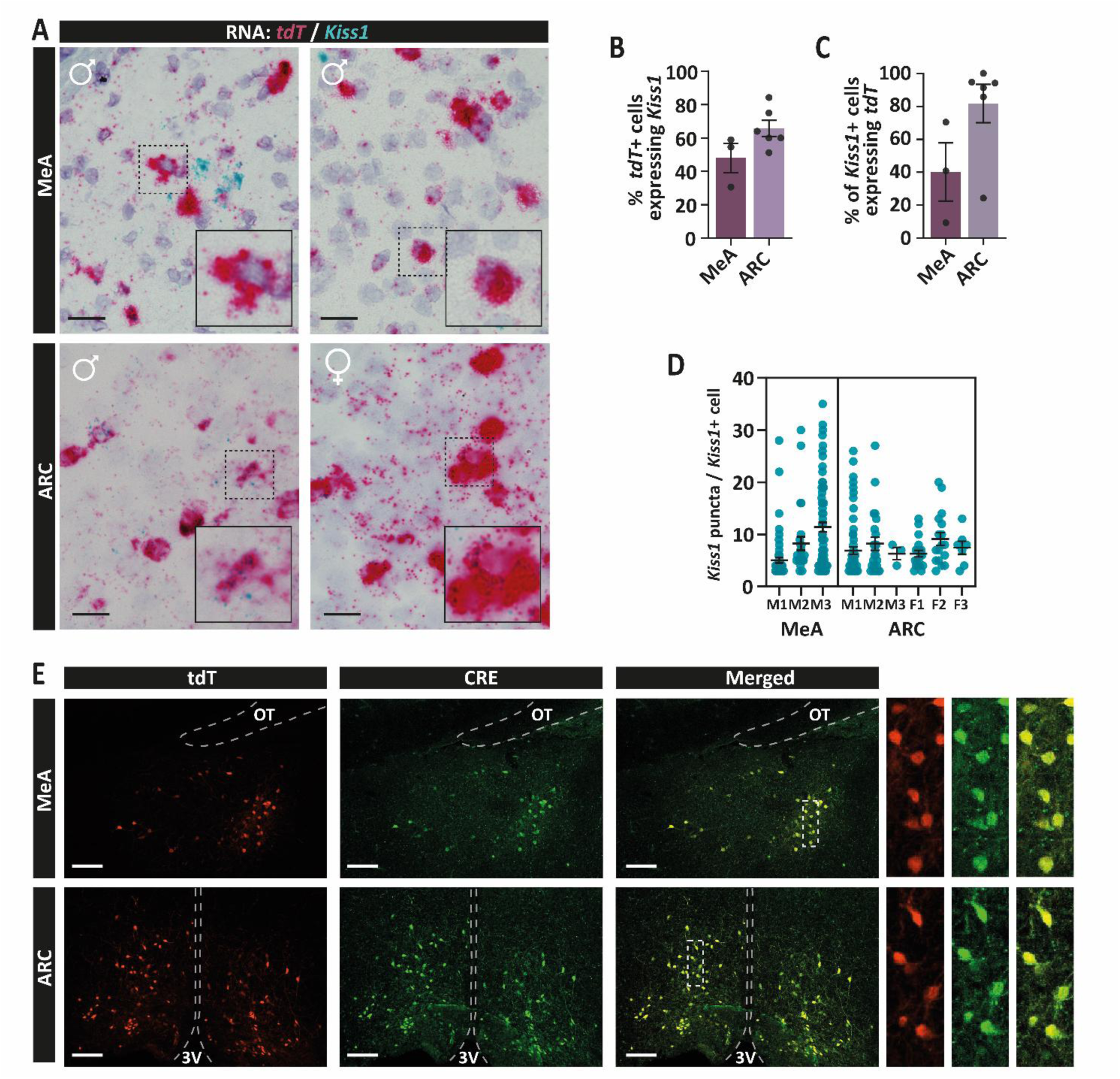
MeA tdTomato positive neurons express the *Kiss1* gene and CRE protein. A. Representative images of *tdT* and *Kiss1* expression in the MeA and ARC (Scale bar = 25µm, inserts are expanded dashed boxes). B. Percentage of *tdT*^+^ cells that had more than 5 *Kiss1* puncta (Welch’s t-test: t(5.62) = 1.26, *p* = 0.2574). C. Percentage of *Kiss1*^+^ cells that were *tdT*^+^ (Mann-Whitney U test: U = 1, *p* = 0.0357). D. *Kiss1* puncta in each *Kiss1*^+^ cell per each mouse (M = Male; F = Female; Nested t-test: t(7) = 0.94, *p* = 0.3799). E. Reporter tdTomato and CRE-antibody labelling in the MeA and ARC (Scale bar = 100µm, on the right expanded dashed boxes). OT = optic tract; 3V = third ventricle.

### Kiss1^MeA^ firing properties differ between sexes and stage of the estrous cycle

Given the age-dependent increase in tdT^+^ cells (Figure 1), mice aged 13-22 wks were used for electrophysiological recordings to maximise the number of neurons available for analysis. Firing properties in *Kiss1*^MeA^ neurons were examined in estrous and diestrous females and in male mice (Figure 3; Table 1). There was no sex- or estrous cycle-dependent differences in input resistance (R_i_), membrane capacitance (C_m_) or resting membrane potential (RMP) (Table 1). To evoke trains of action potentials, *Kiss1*^MeA^ neurons were held at around -70 mV and depolarizing current (5 to 160 pA) was applied for 500 ms (Figure 3A). There was a significant effect of current on the number of spikes but no sex or estrous cycle effect and no interaction (Figure 3B). With increasing current pulses, the increase in firing was significantly slower in estrous females than in males (Figure 3C). There was significantly more spike frequency adaptation (SFA) at the maximum spike number in estrous females compared with males (Figure 3D). All other spike train properties were not significantly different (Table 2). The medium afterhyperpolarization (mAHP) observed at the end of the 500ms spike train was significantly smaller in estrous females compared to males (Figure 3E). In some neurons, hyperpolarization (to around -100 mV) causes an observable voltage sag (Vsag) whereby the membrane initially hyperpolarises and then slowly depolarises (‘sags’) as hyperpolarisation-activated cation channels open. A Vsag was observed in all groups of *Kiss1*^MeA^ neurons, with a significantly smaller amplitude in *Kiss1*^MeA^ of both estrous and diestrous females compared to males (Figure 3F,G). None of the female and 20% of male *Kiss1*^MeA^ neurons fired a rebound action potential at the end of the Vsag (data not shown). Single action potentials (APs) were evoked by 10 ms depolarizing current pulses (Figure 3H). APs fired by *Kiss1*^MeA^ from estrous and diestrous females had a significantly larger amplitude compared to males (Figure 3I). No other differences in AP parameters were seen (Table 1).

**Figure 3.**
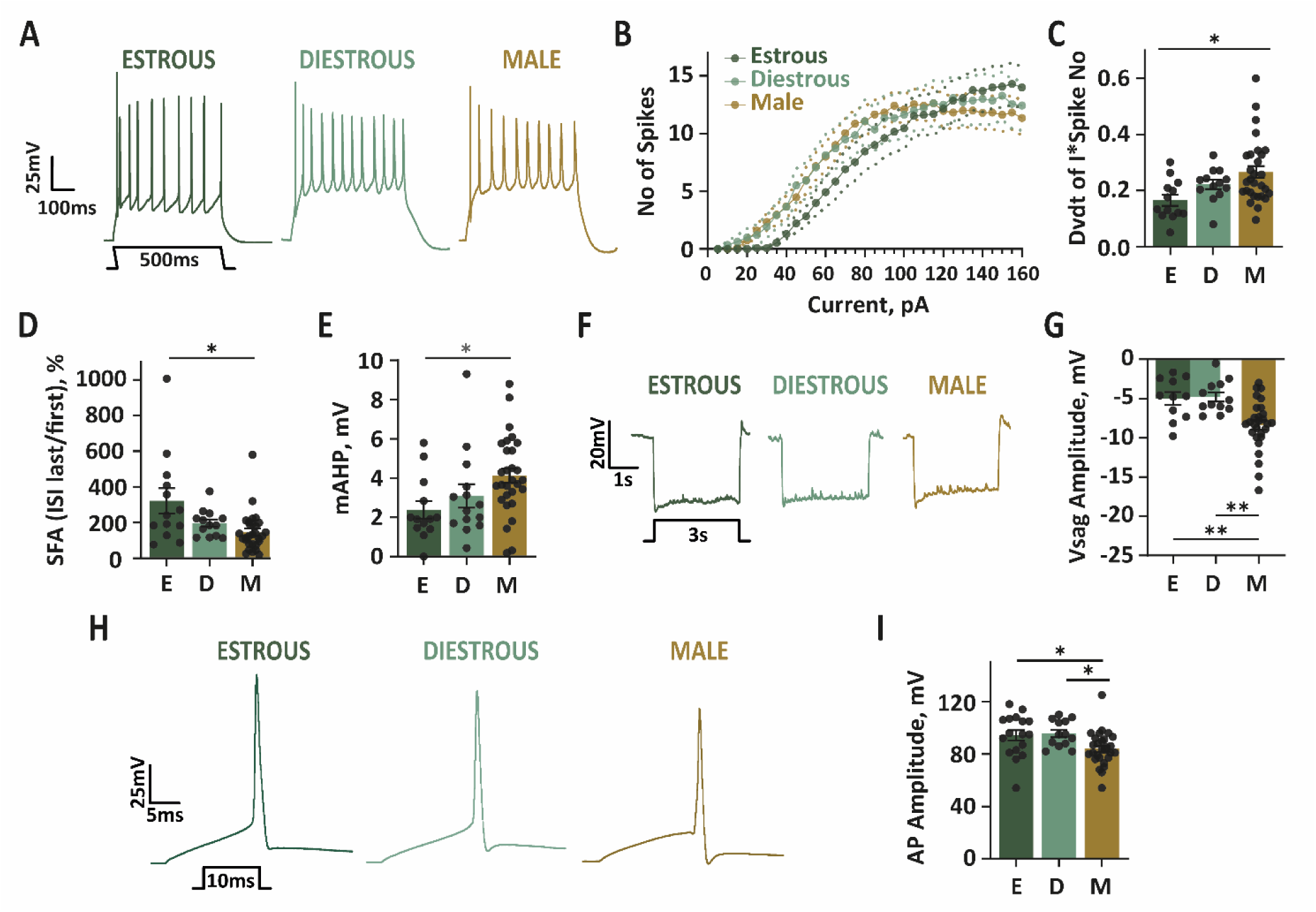
Action potential firing properties of *Kiss1*^MeA^ neurons. A. Representative traces of spike trains occurring in response to 500ms depolarizing current pulses of 80pA from estrous (dark green) and diestrous (light green) females and males (brown). B. Mean number of spikes at each current pulse (Two-way repeated measures ANOVA: Current pulse F(2.226, 118) = 63.12, *p* < 0.0001; Sex/Estrous Stage F(2, 53) = 0.2306, *p* = 0.7948; Interaction F(4.452, 118) = 0.9957, *p* = 0.4181). C. Slope of a linear curve fitted to the 10% minimum to 90% maximum of the input-output curve of the number of spikes fired at each current pulse. (Kruskal-Wallis test: H(2) = 8.464, *p* = 0.0145). D. Spike frequency adaptation (SFA). (Kruskal-Wallis test: H(2) = 9.664, *p* = 0.008). E. Medium afterhyperpolarization (mAHP). (Kruskal-Wallis test: H(2) = 9.244, *p* = 0.0098). F. Representative example traces of voltage sag in response to 3s hyperpolarizing current pulses. G. Amplitude of the depolarizing response (One-way ANOVA: F(2, 48) = 9.301, *p* = 0.0004). H. Representative example traces of single action potentials fired in response to 10 ms depolarizing current pulses. I. Action potential amplitudes measured from threshold to peak. (One-way ANOVA: F(2, 55) = 4.664, *p* = 0.0135). All post hoc P values: \**p* < 0.05; \*\**p* < 0.01

**Table 1.** Electrophysiological properties of *Kiss1*^MeA^ neurons in adult estrous and diestrous female and male mice.

|  | Estrous | Diestrous | Male | P-value |
| --- | --- | --- | --- | --- |
| *Passive Properties* | n = 18 (10) | n = 14 (9) | n = 35 (17) |  |
| RMP, mV | -68.89 ± 1.53 | -66.03 ± 2.82 | -65.58 ± 1.45 | 0.401 |
| R<sub>i</sub>, MΩ | 323.7 ± 22.55 | 429.8 ± 47.72 | 441.9 ± 30.66 | 0.0653 |
| C<sub>M</sub>, pF | 70.7 ± 3.09 | 69.04 ± 5.67 | 72.3 ± 3.79 | 0.8486 |
| *Action Potential Properties* | n = 17 (9) | n = 13 (8) | n = 28 (15) |  |
| Threshold, mV | -47.29 ± 0.71 | -46.54 ± 1.14 | -45.75 ± 0.75 | 0.2809 |
| Amplitude, mV | 94.18 ± 3.94 | 95.62 ± 2.81 | 84 ± 2.47 | **0.0135** |
| WHH, ms | 0.859 ± 0.03 | 0.824 ± .0.03 | 0.858 ± .0.02 | 0.5845 |
| fAHP, mV | -8.748 ± 1.22 | -6.977 ± 2.13 | -5.732 ± 1.21 | 0.3087 |
| Rise Rate, dv/dt | 2.346 ± 0.16 | 2.378 ± 0.14 | 1.972 ± 0.11 | **0.0492** |
| Fall Rate, dv/dt | -1.239 ± 0.06 | -1.324 ± 0.08 | -1.149 ± 0.04 | 0.1255 |
| *Spike Train Properties* | n = 13 (8) | n = 14 (10) | n = 29 (13) |  |
| Max # of Spikes | 15.31 ± 1.5 | 16.43 ± 1.88 | 17 ± 1.12 | 0.7139 |
| Dv/dt of Current*Spike # | 0.165 ± 0.02 | 0.222 ± 0.02 | 0.266 ± 0.02 | **0.0145** |
| Rheobase, pA | 41.15 ± 3.21 | 45 ± 9.27 | 45 ± 4.79 | 0.7629 |
| Frequency at Max Spike #, Hz | 32.43 ± 2.75 | 36.74 ± 3.25 | 35.99 ± 1.94 | 0.5235 |
| First AP Delay, ms | 89.13 ± 15.4 | 98.25 ± 14.1 | 106 ± 14 | 0.5786 |
| mAHP, mV | 2.385 ± 0.45 | 3.094 ± 0.6 | 4.124 ± 0.37 | **0.0098** |
| SFA (ISI last/first)% | 321.8 ± 70.91 | 196.5 ± 20.69 | 147.3 ± 20.24 | **0.008** |
| Depolarization Block | Y: 23.07%<br>N: 76.9% | Y: 42.9%<br>N: 57.1% | Y: 51.7%<br>N: 48.3% | 0.2222 |
| *Voltage Sag Properties* | n = 11 (7) | n = 12 (8) | n = 28 (14) |  |
| Voltage Sag, mV | -4.995 ± 0.81 | -4.81 ± 0.59 | -8.376 ± 0.6 | **0.0004** |

Number of cells, n; number of mice in parentheses.

**Table 2.**
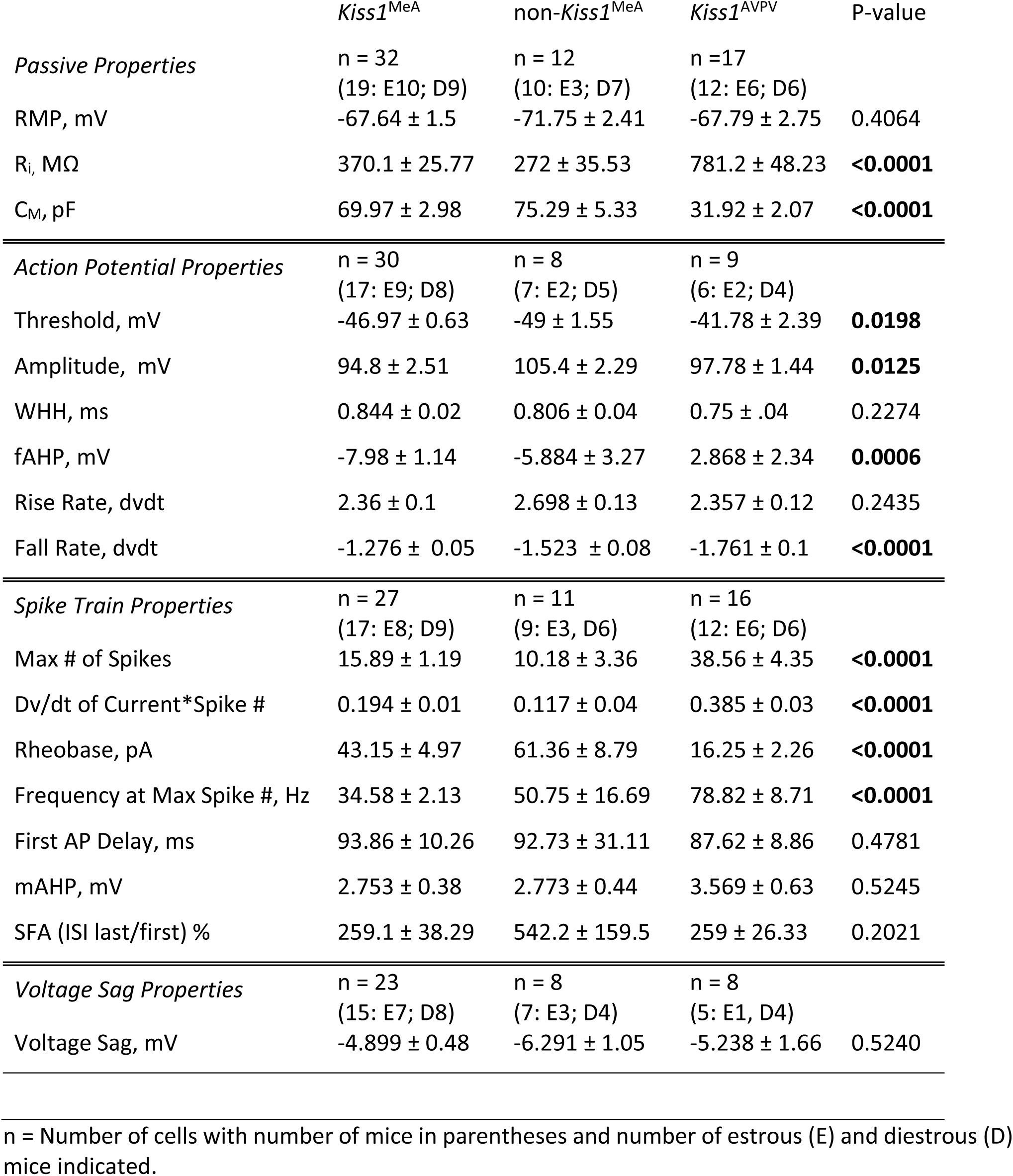
Electrophysiological properties of *Kiss1*^MeA^, non-*Kiss1*^MeA^, and *Kiss1*^AVPV^ in female mice.

Principle Component Analysis (PCA) was performed to examine whether differences in multiple parameters accumulate to a distinct physiological phenotype between *Kiss1*^MeA^ neurons in males and females. Plotting PC1 against PC2 (Figure 4A), as well as PC3 against PC4 (Figure 4B) revealed a general overlap of *Kiss1*^MeA^ neuronal properties from all groups, indicating that there are no distinct electrophysiological phenotypes in males versus females.

**Figure 4.**
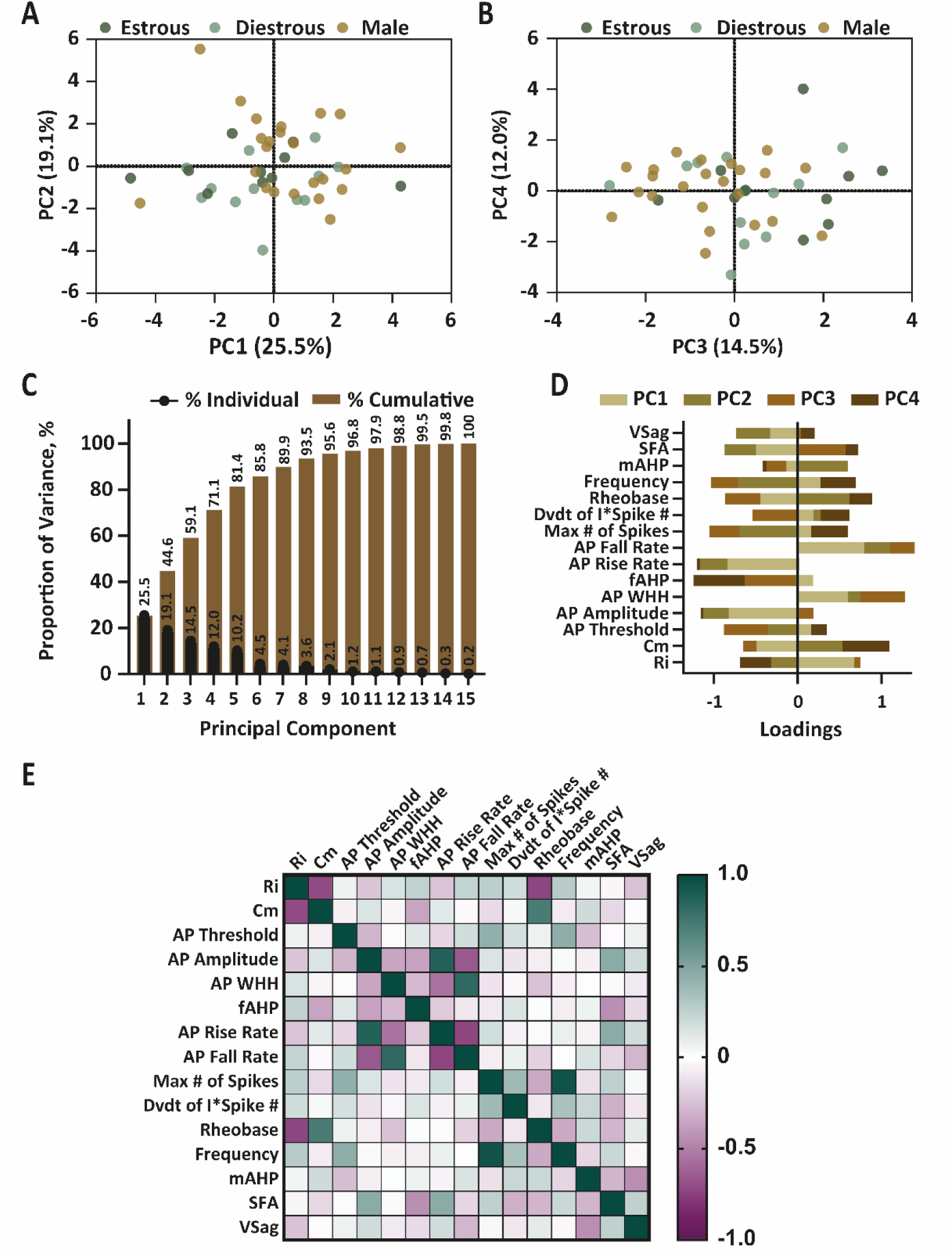
Principal component analysis (PCA) of electrophysiological properties of *Kiss1*^MeA^ neurons did not reveal a distinct physiological phenotype in males versus females. A, B. PC1 and 2 and PC3 and 4 of *Kiss1*^MeA^ in males (n = 23) and estrous (n = 10) and diestrous (n = 12) females. C. Proportion of the explained variance attributed to each principal component. D. Loadings of each variable on each principal component. E. Spearman’s correlation of all variables included in the PCA.

The properties of *Kiss1* neurons at estrous and diestrous were very similar. Each PC is sequentially calculated to explain the variance displayed in the data sets and PCs 1-4 accounted for 71.1% of the cumulative variance (Figure 4C). Loadings of each measured electrophysiological parameter onto any of the four PC’s show no clear bias of any variable to a specific PC, thus no variable could be identified to drive any of the PCs (Figure 4D).

To investigate relationships between physiological properties, a correlation matrix of all variables included in the PCA was generated (Figure 4E). Across groups, capacitance and input resistance were negatively correlated, where higher capacitance and lower input resistance most likely reflects cells with larger cell surface areas. As expected, rheobase, one indicator of neuronal excitability, was positively correlated with capacitance and negatively correlated with input resistance. There was a strong positive correlation between AP rise time and amplitude which indicates that a faster AP rising phase is driving a larger AP amplitude. SFA was positively correlated with both action potential amplitude and rise rate and weakly negatively correlated with the fast and medium AHPs, indicating that, as expected, there is more spike adaptation with fast rising, large APs but less spike adaptation with larger fast and medium AHPs.

### The electrophysiological properties of Kiss1^MeA^ neurons are distinct from Kiss1^AVPV^ neurons and more similar to other neurons in the MeA

*Kiss1*^MeA^ and *Kiss1*^AVPV^ neurons both show sexual dimorphisms: a greater number of *Kiss1*^MeA^ neurons is found in males (Figure 1; Figure S1^12,13,29^) while a greater number of *Kiss1*^AVPV^ neurons is found in females^31^. Both are involved in sexual behaviours. We investigated whether, in female estrous and diestrous mice, the electrophysiological properties of *Kiss1*^MeA^ neurons more closely resemble those of *Kiss1*^AVPV^ neurons or those of other (non-*Kiss1*) MeA neurons. *Kiss1*^MeA^ neurons had a significantly lower input resistance and a larger capacitance than *Kiss1*^AVPV^ neurons but were not different to non-*Kiss1*^MeA^ neurons (Figure 5A, B). RMP did not differ between the three cell types (Figure 5C). When depolarized for 500 ms, *Kiss1*^MeA^ and non-*Kiss1*^MeA^ neurons fired fewer action potentials with increasing current than *Kiss1*^AVPV^ neurons (Figure 5D, E). *Kiss1*^MeA^ and non-*Kiss1*^MeA^ neurons required larger currents to evoke APs (rheobase) and had a slower rate of increasing spike number with each current pulse than *Kiss1*^AVPV^ neurons (Figure 5F, G). The lower maximum number of spikes was accompanied by a lower firing frequency in *Kiss1*^MeA^ and non-*Kiss1*^MeA^ neurons (Figure 5H, I). All parameters are reported in Table 2.

**Figure 5.**
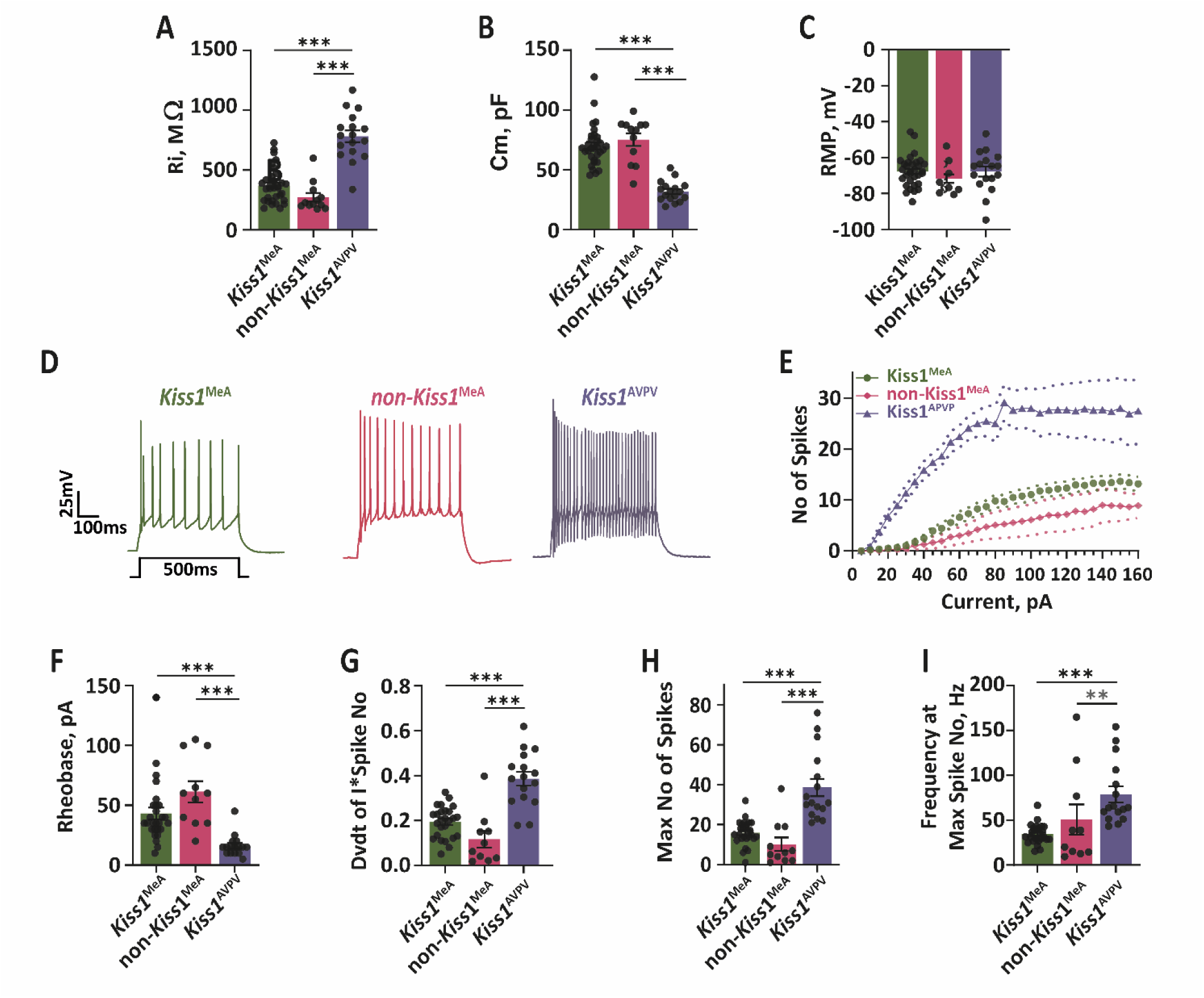
Female *Kiss1* neurons and non-*Kiss1* neurons in the MeA are less excitable than *Kiss1^AVPV^*neurons. A. Input resistance (Kruskal-Wallis test: H(2) = 33.38, *p* < 0.0001). B. Membrane capacitance (Kruskal-Wallis test: H(2) = 35.54, *p* < 0.0001). C. Resting membrane potential (One-way ANOVA: F(2, 58) = 0.9145, *p* = 0.4064). D. Representative example traces of neuronal firing of *Kiss1*^MeA^ (green), non-*Kiss1*^MeA^ (pink), and *Kiss1*^AVPV^ (purple) neurons from female mice in response to a 500ms depolarizing current (80pA) pulse. E. The mean number of spikes at each increasing current pulse (Two-way repeated measures ANOVA: Interaction F(3.106, 79.21) = 3.818, *p* = 0.0122). F. Current pulse at which the first action potential was fired (Kruskal-Wallis test: H(2) = 26.75, *p* < 0.0001). G. Slope of the linear section of each current pulse*number of spikes curve (Kruskal-Wallis test: H(2) = 26.15, *p* < 0.0001). H. Maximum number of Spikes fired (regardless of current pulse amplitude) (Kruskal-Wallis test: H(2) = 30.64, *p* < 0.0001. I. Firing frequency at the maximum spike number (Kruskal-Wallis test: H(2) = 21.94, *p* < 0.0001). All post hoc *p* values: \**p* < 0.05; \*\**p* < 0.01; \*\*\**p* < 0.001.

To visualize combined differences between *Kiss1*^MeA^, non*-Kiss1*^MeA^, and *Kiss1*^AVPV^ neurons, PCA was performed. To preserve a reasonable n-number per cell-type, only passive properties and spike train firing data were included in this analysis. PC1 and PC2 show a visible shift in *Kiss1*^MeA^ and *Kiss1*^AVPV^ neuronal properties with only few *Kiss1* neurons from either brain area occupying a similar plot space on the PC1 axis (Figure 6A). Non-*Kiss1*^MeA^ neurons largely occupy a similar space to *Kiss1*^MeA^ neurons. PC1 and PC2 account for 70.8% of the cumulative variance, with 55.9% of the variance being accounted for by PC1 (Figure 6B). To investigate whether this is reflected in the proportion of variables loading into PC1 the variable loadings for PC1 and PC22 were plotted. Loadings for PC1 were predominantly influenced by Ri, Cm, the maximum number of spikes, rheobase, frequency of firing, and the rate at which spike numbers increase with each current pulse (Figure 6C). Only SFA and the mAHP predominantly influenced loadings for PC2. The variables loading PC1 were highly correlated with each other while those loading PC2 were not (Figure 6D).

**Figure 6.**
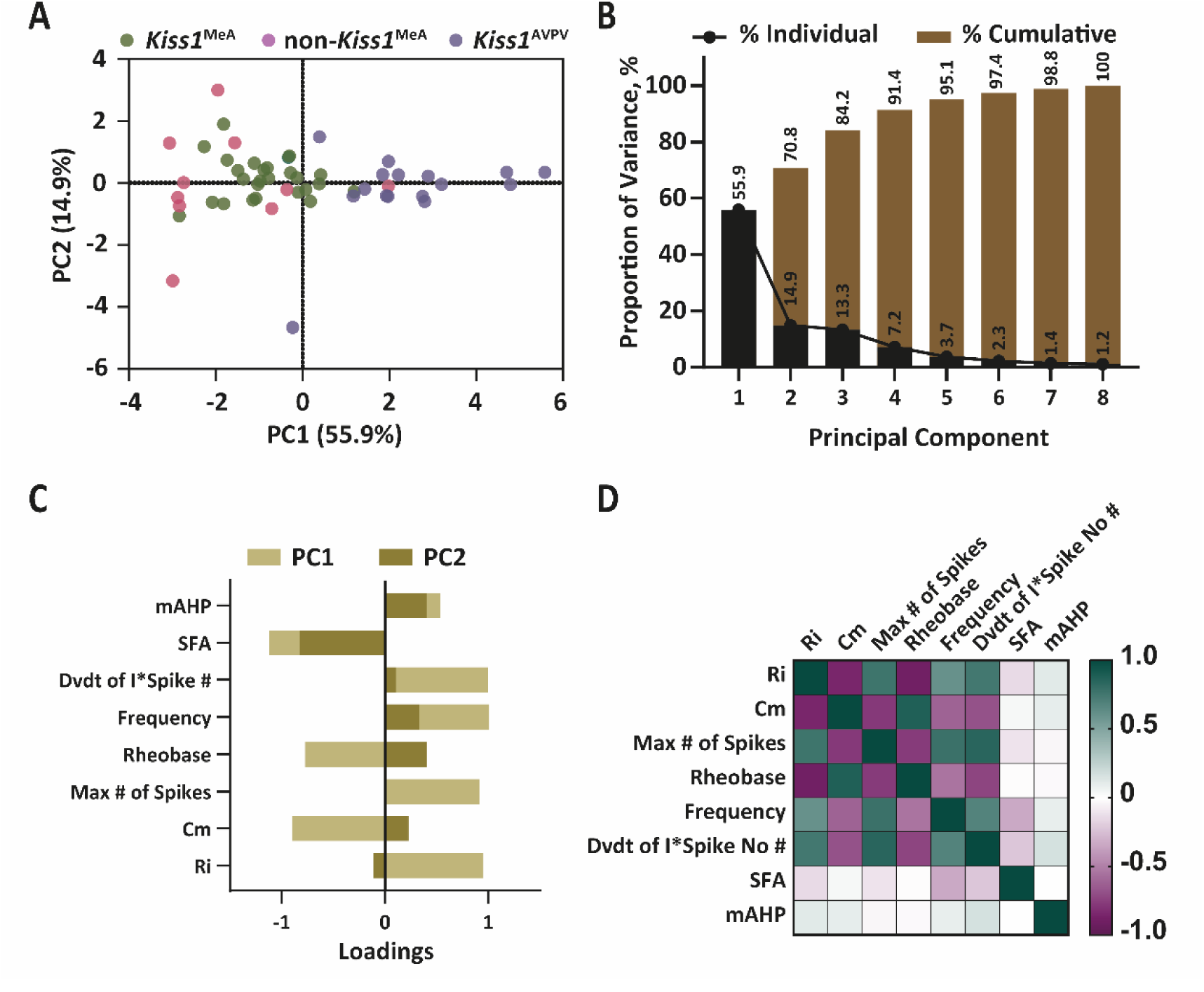
PCA reveals two electrophysiologically distinct kisspeptin populations in the MeA and AVPV. A. PC1 and 2 of firing properties from *Kiss1*^MeA^ (n = 27), non-*Kiss1*^MeA^ (n = 10), and *Kiss1*^AVPV^ (n = 14) in female mice. B. Proportion of the explained variance attributed to each principal component. C. Loadings of each variable on each principal component. D. Spearman’s correlation of all variables included in the PCA.

Hypothalamic *Kiss1* neurons must fire at frequencies greater than 10Hz to release kisspeptin onto GnRH neurons and evoke pulsatile GnRH and LH release^34,46,47^. The amount of current required to evoke sustained firing (>400ms) at 10Hz was compared between *Kiss1*^MeA^ and *Kiss1*^AVPV^ neurons. The threshold current for evoking firing at frequencies greater than 10Hz was significantly smaller in *Kiss1*^AVPV^ neurons than in *Kiss1*^MeA^ neurons, indicating that *Kiss1*^AVPV^ neurons can more readily achieve action potential firing frequencies that favour neuropeptide release (Figure 7).

**Figure 7.**
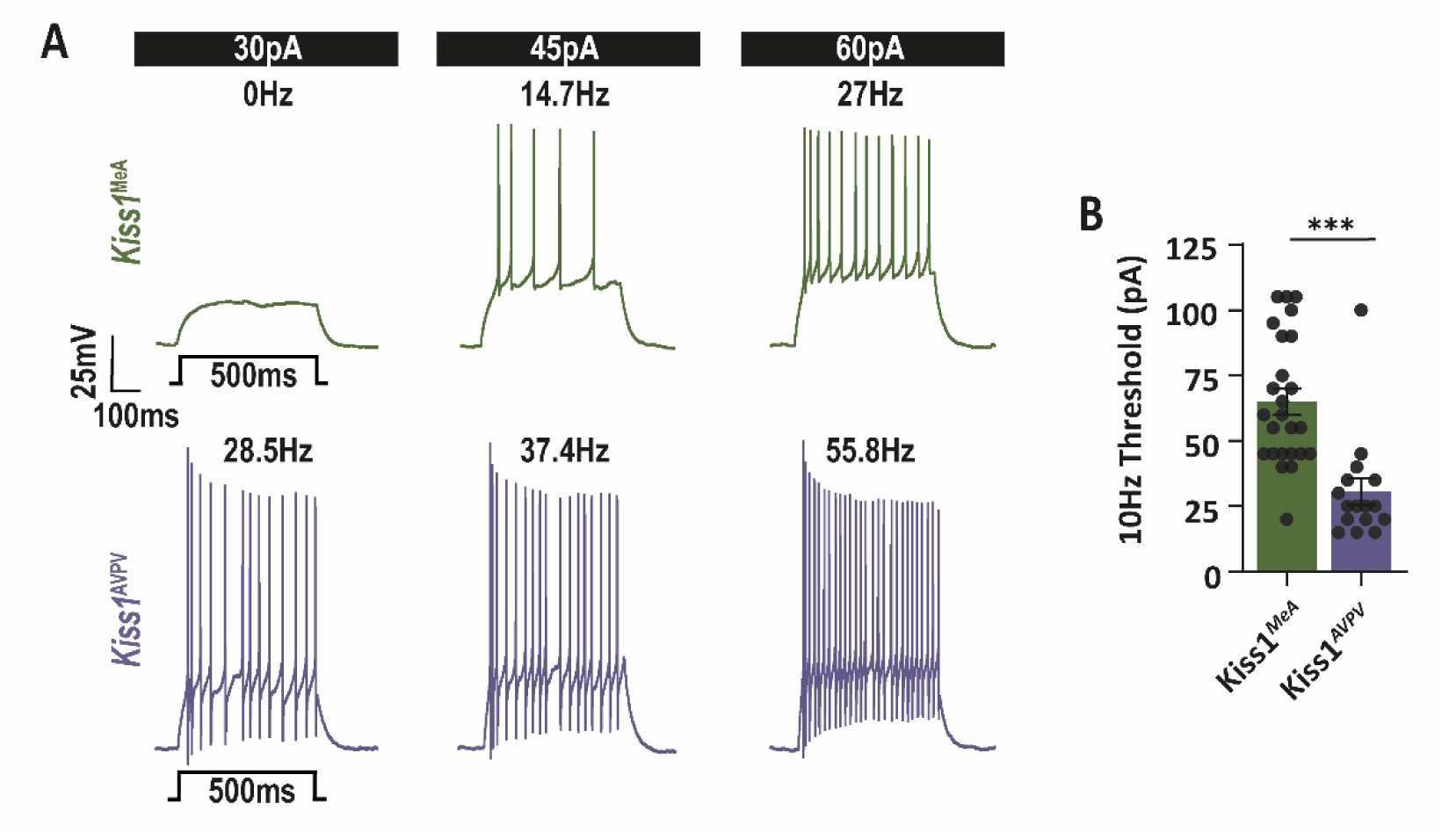
*Kiss1*^MeA^ neurons require larger current input to fire at 10Hz than *Kiss1*^AVPV^. (A) Representative traces of *Kiss1* neurons in the MeA and AVPV at 30pA, 45pA, and 60pA 500ms current steps. Firing frequency of displayed traces indicated above. (B) Lowest current at which a neuron fired at a minimum of 10Hz for a sustained period (400ms) of the 500ms current step (Mann Whitney U = 38, *p* < 0.0001). \*\*\**p* < 0.001

## Discussion

*Kiss1* neurons are modulated by sex steroids, providing important gonadal physiological feedback to brain circuits involved in reproduction and reproduction-related behaviours. Here, the neuronal firing properties of *Kiss1*^MeA^ neurons were found to be sexually dimorphic, with differences most pronounced between estrous females and males. Additionally, the firing properties of *Kiss1*^MeA^ neurons differed from the hypothalamic *Kiss1*^AVPV^ population, suggesting a unique *Kiss1* neuronal cell type with different output activity in the MeA.

The reported sexual dimorphism in *Kiss1*^MeA^ neuron number was confirmed by counting the number of tdTomato neurons labelled through expression of the CRE recombinase from the *Kiss1* promoter, providing the first quantification across a range of adult ages. Males had ∼6x as many tdT^+^ cells as females at the oldest time point studied (20wks). However, compared to the *Kiss1^Cre-v2-tdT^* mouse line, the number of tdT^+^ neurons in *Kiss1^Cre-tdT^* mice was lower at all time points measured. The lower numbers of MeA tdT^+^ cells in *Kiss1^Cre-tdT^* mice could result from an under-estimate, due to undetectable expression of tdTomato in some *Kiss1* neurons, although our cell counts are consistent with *Kiss1* expression in the MeA reported by *in situ* hybridisation^12,13,40^. RNAscope confirmed that, in the MeA of *Kiss1^Cre-tdT^* mice, ∼50% of *tdT*^+^ cells also expressed detectable *Kiss1*, similar to ∼60% in the ARC. Co-localisation of kisspeptin immunoreactivity with tdT in the ARC has previously been reported at 81-90% in GDX males and females of this mouse line^15^. Lower numbers observed here might be accounted for by lower *Kiss1* levels in the ARC of intact mice, where negative estrogen feedback is active. *Kiss1* expression in the MeA is known to be lower than in the ARC^19,48^, and thus might be underestimated here due to limits of detection of the RNAscope assay. We also found that some *Kiss1*^+^ neurons in the MeA (and, to a lesser extent, in the ARC) did not express detectable *tdTomato* transcripts. It is not known why *Kiss1* promoter activity is not driving tdTomato expression in these cells. Given that 87% of tdT^+^ cells were immunoreactive for CRE, this suggests that the CRE levels might not be high enough to activate the tdTomato reporter. It is important to note that both immunohistochemistry for the kisspeptin protein and *Kiss1* RNAscope are notoriously difficult to achieve in the MeA due to the low levels of RNA and protein expression^37^.

The age dependent increase in the number of tdT^+^ neurons was observed in both mouse lines and is consistent with data from a *Kiss1^Cre^*rat line, where a notable increase in tdTomato labelling in the amygdala was observed between 7 and 10 weeks^29^. Both the rat and the mouse models rely on visualisation with a tdTomato reporter line, meaning that after even transient *Kiss1* expression, the tdTomato signal is permanently switched on. Thus, an increase in tdT^+^ cells in adulthood could indicate either a temporal increase in cells expressing *Kiss1* or a system in which the overall number of cells expressing *Kiss1* is consistent but different neurons express *Kiss1* at different times. Interestingly, the number of tdT^+^ neurons in the mouse MeA, especially in the *Kiss1^Cre-tdT^* line, was relatively low immediately after puberty, when mice can mate; this was surprising, given the role of *Kiss1*^MeA^ neurons in sexual behaviour. TdT^+^ cell counts did not support the idea that *Kiss1* neuron expression increases with sexual experience.

The higher number of tdT^+^ cells further into adulthood informed a pragmatic decision to perform electrophysiological experiments in mice at 13-20 wks of age. This is the first report of *Kiss1*^MeA^ neuron electrophysiological properties. *Kiss1*^MeA^ neurons in estrous females showed a slower change in excitability (measured as the rate of change in the number of action potentials with increasing current pulses) compared to males and also showed more SFA (slowing of firing frequency during sustained depolarizing current); this could be due to lower excitability, but might also be accounted for by reduced expression of ion channels that allow sustained high frequency firing, such as calcium-activated potassium and chloride channels driving the mAHP and fAHP^49,50^. After-hyperpolarisations remove the inactivation of voltage-gated sodium channels, making them available for recruitment and supporting higher frequency firing. Consistent with this, the mAHP was significantly smaller in *Kiss1*^MeA^ neurons from estrous females compared with males and SFA was weakly inversely correlated with mAHP size.

Principal component analysis (PCA) confirmed that *Kiss1*^MeA^ neurons show similar firing properties between sexes. Neither individual measurements nor PCA showed cycle-dependent differences. We defined estrous and diestrous within a tight time window (see Methods). This should mean that during estrous tissue was collected shortly after the LH surge and after the high estrogen levels during proestrous^51^. However, the estrous cycle in mice cannot be assumed to function in strict 24 hour time blocks. Furthermore, there are differences in reported sex steroid levels across the cycle and there may be differences across mouse strains^51–53^. Naturally occurring variability may be masking differences in *Kiss1^MeA^* firing properties during the estrous cycle. Nonetheless, our observation of limited cycle-dependence of *Kiss1*^MeA^ neuronal properties is consistent with RNAseq data^37^ under controlled sex steroid levels: *Kiss1*^MeA^ neurons in ovariectomized females mice with and without estrogen replacement showed only 45 differentially expressed genes, suggesting that modulation of gene expression during the cycle is moderate.

Hypothalamic *Kiss1* neurons are very effective modulators of GnRH neuron activity, and thus LH and FSH release from the pituitary^1–6^. Both AVPV and ARC *Kiss1* neurons generate high frequency action potential firing^36^. *Kiss1*^MeA^ neurons also modulate LH pulses in both females and males via either direct or indirect projections to GnRH neurons^21,25,54^. Vesicular neuropeptide release requires high frequency firing, ideally burst firing patterns^55^; indeed it has been shown that low frequency activation of AVPV neurons predominantly causes release of amino acid neurotransmitters (GABA and glutamate) that are co-expressed by hypothalamic *Kiss1* neurons, rather than kisspeptin release, leading to short term activation of *GnRH*^POA^ neurons^46^. By contrast, high frequency activation stimulates kisspeptin release, which causes a delayed but longer lasting increase in *GnRH*^POA^ neuron firing frequency^56^. The firing properties of *Kiss1*^MeA^ neurons were therefore compared with *Kiss1*^AVPV^ neurons. *Kiss1*^MeA^ neurons were larger and had a lower input resistance, distinct firing patterns and lower excitability than *Kiss1*^AVPV^ neurons. This raises the question of whether *Kiss1*^MeA^ neuronal stimulation of *GnRH*^POA^ neurons (and thus LH release) involves kisspeptin release, or GABA and/or glutamate release. Transcripts for GABA and glutamate production have been identified in bulk RNAseq data from female *Kiss1*^MeA^ neurons and RNAscope data in male mice and indicates that ∼70% of *Kiss1*^MeA^ neurons express the GABAergic marker *Vgat* while 30% express the glutamatergic marker *Vglut*, with a few cells expressing both^24,37^. Our findings, of much lower firing frequencies of *Kiss1*^MeA^ neurons, would suggest that they might preferentially release GABA or glutamate rather than kisspeptin: in the hypothalamus, locally within the MeA, and in other target brain regions. The consequences of the lower firing frequency of *Kiss1*^MeA^ neurons in terms of neurotransmitter versus neuropeptide release require further investigation.

*Kiss1*^MeA^ neurons showed more similarities to other (non-*Kiss1*) MeA neurons, although the PCA revealed that they are not completely overlapping in their electrophysiological properties. This is likely because neurons in the MeA are highly diverse, with local inhibitory neurons and different types of projection neurons contrasting in their firing patterns^57–59^. Keshavarzi et al. (2014) identified two types of non-GABAergic and three types of GABAergic neurons based on their electrophysiological firing properties. *Kiss1*^MeA^ neuronal firing properties seem to be most similar to multipolar type II non-GABAergic neurons^60^.

In conclusion, the electrophysiological properties of *Kiss1*^MeA^ neurons are reported for the first time, showing that their firing properties differ between male mice and estrous female mice. Interestingly, they have unique electrophysiological properties to kisspeptin neurons in the AVPV of the hypothalamus, showing lower excitability and action potential firing frequency. This might have consequences for the release of kisspeptin versus amino acid co-transmitters in hypothalamic, local amygdala and other brain targets.

