## Supplemental Figure 1 for "*Kiss1* neurons in the medial amygdala of mice are sexually dimorphic and unique from hypothalamic anteroventral periventricular *Kiss1* neurons in their action potential firing properties"

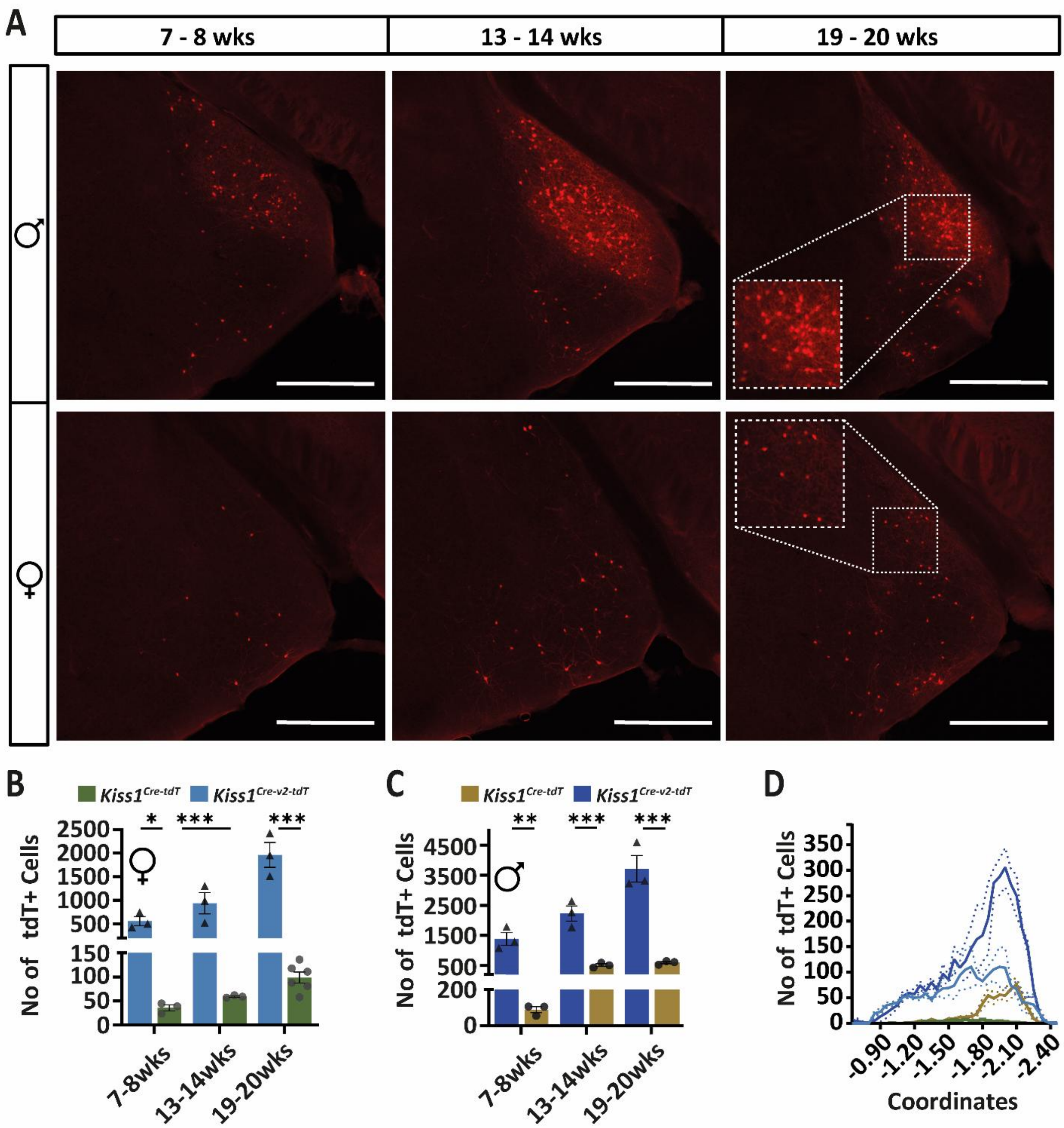


**Supplementary Figure 1. More tdT labelling in the MeA is the _Kiss1_$^{Cre\text{-}v2\text{-}TdT}$ transgenic mouse line.** A. Representative images of tdT$^+$ cell distribution in 50µm slices in the _Kiss1_$^{Cre\text{-}v2\text{-}TdT}$ mouse line [41]. B. Number of tdT$^+$ cells at 7+8, 10+11, 13+14, and 19+20wks old in female and male _Kiss1_$^{Cre\text{-}v2\text{-}TdT}$ and _Kiss1_$^{Cre\text{-}tdT}$ mice. C Mean anterior to posterior distribution of tdT$^+$ cells in both mouse models at 19+20wks. All post hoc P values: *$p < 0.05$; **$p < 0.01$; ***$p < 0.001$. (Scale bar = 500µm). Data from 3 mice in all cases.
