## Supplemental Figure 2 for "*Kiss1* neurons in the medial amygdala of mice are sexually dimorphic and unique from hypothalamic anteroventral periventricular *Kiss1* neurons in their action potential firing properties"

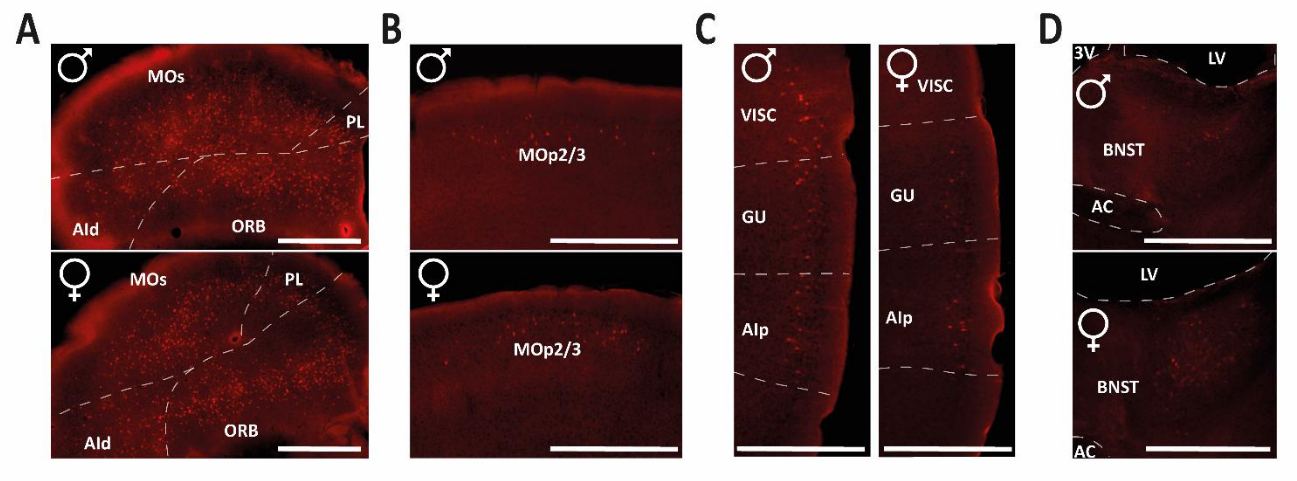


**Supplementary Figure 2. Ectopic tdT labelling in the _Kiss1_$^{Cre\text{-}v2\text{-}TdT}$ transgenic mouse line**. Example images from one male and one female mouse at 19 weeks old in anterior to posterior order. A. Anterior cortex, including secondary motor area (MOs), prelimbic area (PL), orbital area (ORB), and the dorsal agranular insular area (AId). B. Primary motor area layer2/3. C. Cortex, including visceral area (VISC), gustatory area (GU), and the posterior agranular insular area (AIp). D. Bed nucleus of the stria terminalis (BNST). (Scale bar = 600µm).
